# Cortical representation variability aligns with in-class variances and can help one-shot learning

**DOI:** 10.1101/2021.01.27.428518

**Authors:** Jiaqi Shang, Eric Shea-Brown, Stefan Mihalas

## Abstract

Learning invariance across a set of transformations is an important step in mapping high-dimensional inputs to a limited number of classes. After understanding the set of\ invariances, can a new class be learned from one element? We propose a representation which can facilitate such learning: if the variability in representing individual elements across trials aligns with the variability among different elements in a class, then class boundaries learned from the variable representations of one element should be representative of the entire class. In this study, we test whether such a representation occurs in mouse visual systems. We use Neuropixels probes recording single unit activity in mice observing 200 repeats of natural movies taken from a set of 9 continuous clips. We observe that the trial-by-trial variability in the representation of individual frames is well aligned to the variability in representation of multiple frames from the same clip, but not well aligned to the variability among frames from different clips. Thus, the variable representations of images in the mouse cortex can be efficiently used to classify images into their clips. We compare these representations to those in artificial neural networks. We find that, when introducing noise in networks trained for classification (both feed-forward and recurrent networks), the variability in the representation of elements aligns with the in-class variance. The networks which best reproduce the in-vivo observed directions of variability were those trained on a hierarchical classification task. Taken together, these results point to a solution which the cortex can use for one-shot learning of a class: by using noise as a mechanism for generalization. This is a potential computational explanation for the high level of noise observed in the cortex.

## Introduction

A salient feature of the neuronal activity in the mammalian cortex is the surprising level of trial-by-trial variability (Tolhurst et al. 1983; Shadlen and Newsome 1998; Faisal et al. 2008), which has been has been quantified using large scale recordings (de Vries et al. 2020). Generally, the variability is viewed as a nuisance, and multiple experimental (Newsome, Britten, and Movshon 1989) and theoretical (Averbeck, Latham, and Pouget 2006; Abbott and Dayan 1999) analyses have been performed looking at how variability limits the accuracy of coding. Recent work has explored the limitations caused by variability in coding of minute feature differences (Rumyantsev et al. 2020). An implicit assumption underlying these analyses is that the function of the cortex is to represent the visual stimuli as accurately as possible.

Other studies look at noise as a feature, not a bug. For example, (Echeveste and Lengyel 2018) discuss the role noise can play in inference. Using inspiration from the role of noise in helping artificial neural networks generalize (Srivastava et al. 2014), we have proposed that the role of the noise in the cortex is also to help generalization (Hu et al. 2019). In this previous study we explored the distribution of the trial-by-trail variability and found it better fit by gaussian mixture models (which are similar to those observed in networks trained with dropout) than normal distributions. We have also looked at the direction of the variability, and, for a few example mice, we found an alignment between the variability in representations of a frame in a movie and the variability of responses within a continuous clip (Hu et al. 2019).

In the current study, we extend this alignment result, focusing on the role it can play in classification and one-shot learning. Here, we characterize this alignment mathematically by describing the variance of representation of different examples in a class in feature space (*in class variance*). Separately, we look for the variation of the representation to multiple repeats of one example when noise is injected (*exemplar variance*). When comparing the variance of one exemplar, we can compare it to the variance between other exemplars in the same class (*in same class variance*), or in a different class (*other class variance*). Similarly, we can describe the alignment to the direction orthogonal to the class boundary for a linear classifier (*between class axis*). We construct a new measure to quantify the alignment of the subspace spanned by noisy representation of an example, the subspace spanned by classes of examples, and the best linear separator between these classes (see Figure 1 and the methods section). We show significant alignment of the variability to the exemplar and in-class variance. This alignment is deleterious for stimulus identification but useful for low or one-shot learning in which a new class boundary can be learned from a small number of or a single example of each class. We compare such an alignment with feedforward and recurrent artificial neural networks trained for image classification. In both cases, we find that these trained networks, when responding to images and unstructured input noise, develop a similar alignment. The most similar alignment to that in cortical recordings is observed when for artificial networks trained to perform hierarchical classification.

**Figure 1:**
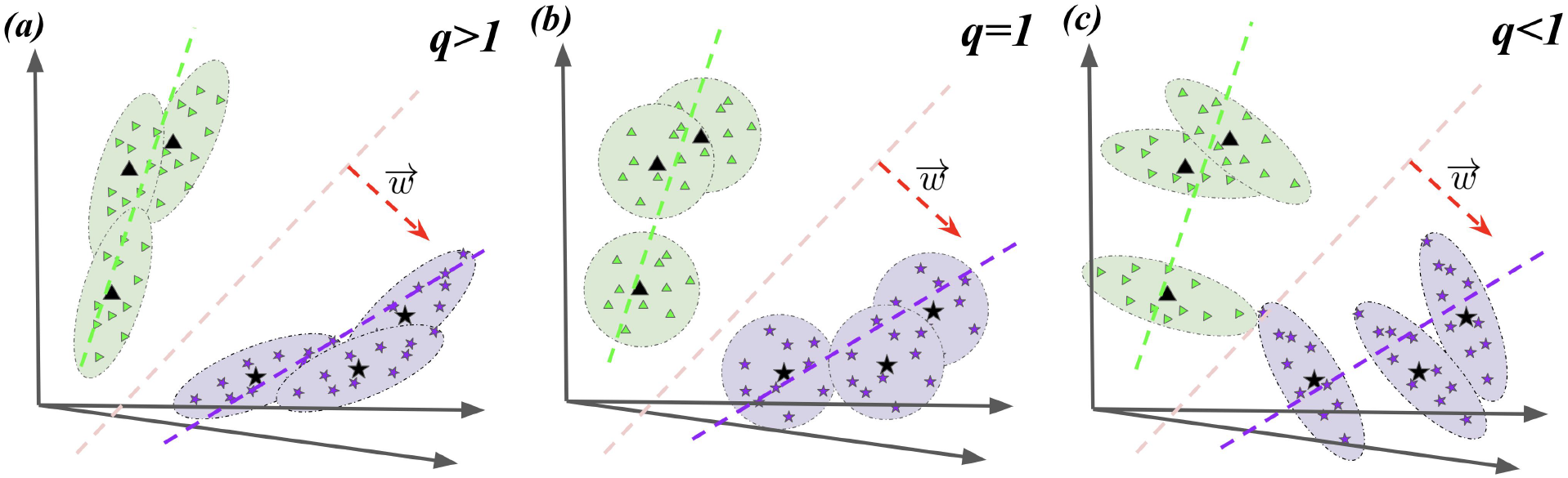
Different possible structures of exemplar variance. In all diagrams, the two classes are color-coded with green and purple. The trial-averaged representation for each stimulus is shown with black triangles/stars, with the exemplar variance shown with green/purple ellipses. The in-class variance is shown in dashed green/purple lines and the between class axis is shown in dashed red line. The exemplar variance can be aligned with (Fig a, q>1), independent from (Fig b, q=1) or orthogonal (Fig c, q<1) to the in-class variance.

The number of possible stimuli which can impede on a mammalian retina is enormous. Even for the relatively small number of receptors in mice (Jeon et al. 1998)), the number of possible stimuli is larger than the number of atoms in the universe. Faced with this, it is natural that part of the function of the visual cortex is to perform a many-level hierarchical classification.

Hierarchical classification is a common problem, where the goal is to differentiate stimuli at different levels: e.g. to differentiate between visual images of two cats from dogs, and produce an output both at the level of the larger class (is it a cat or dog?) and at the subclass (which of the individual cats is in the image?).

Given 2 classes of stimuli, the variability of trial-to-trial representations of stimuli within classes can take different structures in relation to the trial-averaged representation across stimuli of the class. Specifically, this trail-to-trial variability could align with, be unrelated to, or orthogonal to the within-class variation structure (Figure 1). The potential variability structures can be quantified by q, defined as the fraction of variance of the exemplar variance explained along the direction of the in-class variance, normalized by the fraction of variance explained along any random direction. Thus, exemplar variance aligned to in-class variance will have a q larger than 1, while exemplar variance orthogonal to in-class variance will lead to a q smaller than 1.

We are interested in looking at the signatures of such alignment in neural data. While possible, it is difficult to train rodents in object classification tasks (Zoccolan et al. 2009). To allow analysis of large scale recordings in mice, we make an assumption that images in natural movies which are close in time and can smoothly transform from one to another belong to the same class. It is a natural assumption for learning classes, as the amount of examples available for such an unsupervised learning vastly outnumber possible supervised examples, even for humans, and the cases in which objects smoothly change identities are rare. An unsupervised learning to map nearby frames in a movie to nearby representations has been shown to create representations similar to those observed in mammalian cortices (Zhuang et al. 2020) and similar to those using supervised training (Yamins et al. 2014). The standard set of stimuli in the Allen Brain Observatory (de Vries et al. 2020) does not contain natural movies with multiple statistics, and we used follow-up recordings (Hu et al. 2019) in which mice are passively viewing highly repeated stimuli composed of multiple short clips with varied statistics.

We find that such an alignment can have an important role in one-shot learning. Using the cortical recordings, we added a single exemplar of each of the last two classes. When this exemplar is perturbed with noise, we demonstrate that the capacity of the learned classification boundary from one example to generalize is significantly dependent on the alignment of the noise added to the in-class variance. In particular, alignment with q>>1 -- as observed in the cortical data -- leads to substantially improved generalization. Thus we conclude that the observed structure of the noise in the cortex can enhance one-shot learning of classification boundaries, in cases -- as for the movie clips studied here -- when the directions of class variance can be generalized from previously learned classes.

## Results

### Single-trial variability in mouse V1 aligns with in-class variance

Different natural movie clips are presented to mice for 200 trials, and the neural responses in multiple visual areas are recorded with Neuropixels probes (Methods I). We define a stimulus class to be the set of frames from the same movie clip. The neural response to the same frame varies across trials, defining the exemplar variance. The variability in the mean across trials of different stimuli in a class is the in-class variance. The plane which best differentiates two classes, characterized by the direction normal to the plane, is defined as the between-class axis. The alignments between the exemplar variance of stimulus to its same/other in-class variances as well as between-class axis are quantified by the alignment metric q (for details, Methods, II).

In more detail, same/other in-class variance is represented by the top principal component (PC) of the trial-averaged response to stimuli of the same/other class. Alignment analysis shows that the fraction of exemplar variance along the same/other class variance is significantly larger than chance (Figure 2a top, signed-rank test p value <2.2e-16), given by values of q>1. We also quantify the alignment of the exemplar variance along directions that differentiate between the class of the stimulus and another, different class. Specifically, the classification axis is defined as the linear SVM classifier defined on the trial-averaged representations for stimuli of the two classes. While significantly less aligned than for the same in-class variance, the exemplar variance is still more aligned to the classification axis than chance (Figure 1a bottom, rank sum test p value <2.2e-16).

**Figure 2:**
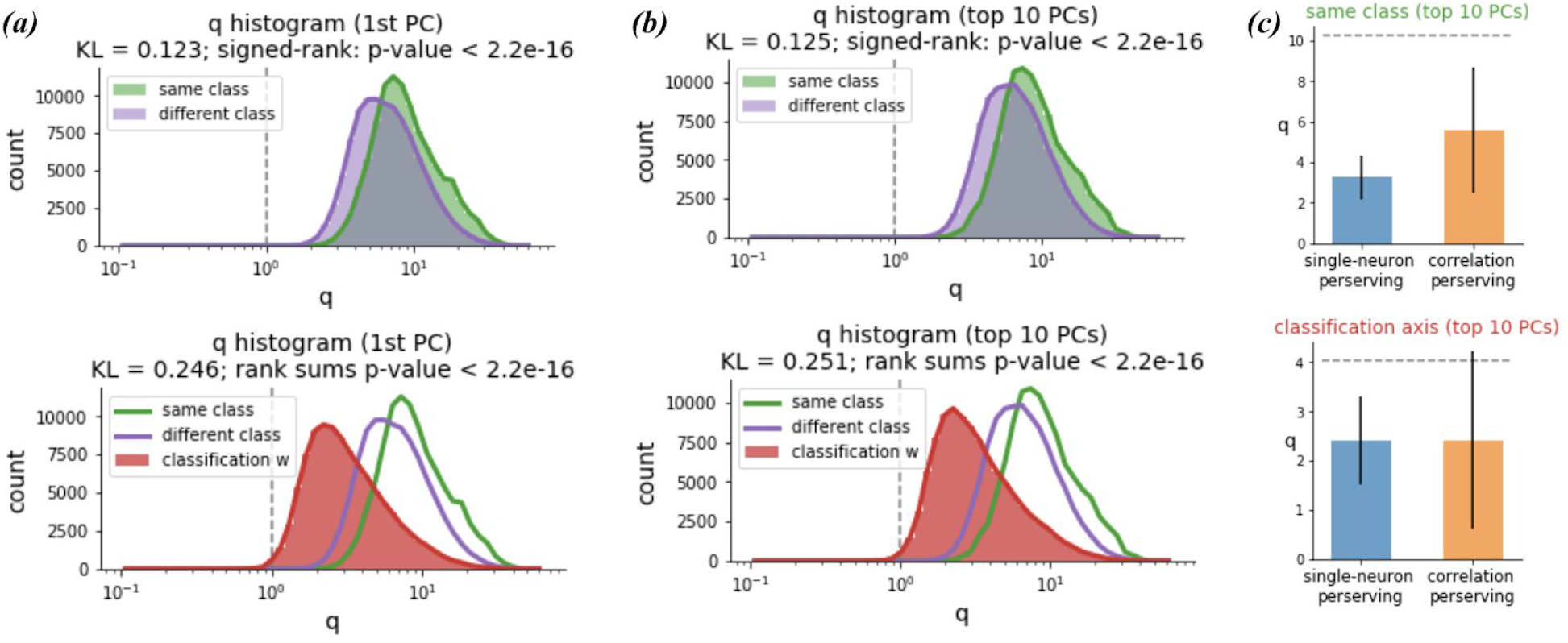
Variability alignment in the neural data. a) q histogram of variability alignment to the same/different class variance (top) and to the between-class axis (bottom). b) q histogram of the variability alignment, with the top 10 PC components included to account for the multi-dimensional population structure. c) single-neuron and population-level structure contribution. (n=7 mice)

So far, the same/other variance and classification structure are represented by a single axis in the high dimensional neural space. Such simplification brings the potential concern that the structures may span multiple axes and the exemplar variance may be aligned to axes other than the single axis included in the analysis. To account for multiple axes in the q alignment quantification, we repeat the analysis with the top 10 PCs of the class trial-averaged response as representations of the same/other class variance structure, weighted by their corresponding explained variance ratio (methods II). The averaged exemplar variance along the same/other class variance across 10 PCs remains significantly larger than chance (Figure 1b top, signed-rank test p=0.0). Similarly, when taking the top 10 classification axes into account (methods II), the exemplar variance remains aligned to the classification axes, while significantly less aligned compared to the same/other in-class variance (Figure 1b bottom, rank sum test p value <2.2e-16).

Next, we ask how much of the alignment is due to single-neuron properties (neurons with larger trial-averaged activity also tend to have larger variance) or the population-level structure (correlations between pairs of neurons). To test for the distribution of single-neuron properties, we retain the empirical distribution and disrupt any population-level structure through trial-shuffling. The simulated data that maintains the single-neuron properties recover 37.4 ± 13.6% of the alignment (Figure 2c). To test for the role of the population-level structure, we retain the pairwise neural correlations and break the single-neuron properties by both changing the single-neuron variances and replacing the single-neuron distributions with a “null’’ gaussian description. Specifically, we simulate from a multivariate Gaussian distribution drawn from a covariance matrix with the same off-diagonal terms as the original data, but with diagonal terms shuffled. The simulated data recover 55.7 ± 6.6% of the alignment, larger than the portion recovered by the single-neuron properties. In summary, both single-neuron and population level statistics contribute to the observed alignment of exemplar variance with the same in-class variance.

### Trained, but not untrained, neural networks exhibit the same general trends in variability alignment

To probe the origin of the aligned variability that is observed in the neural data, we use a convolutional neural network as a model for the visual system. Specifically, we use the Alexnet (Krizhevsky et al. 2012)network, trained on the CIFAR10 dataset (Krizhevsky 2009) (Methods III). To introduce variability in the network response, we pass in the inputs with added Gaussian noise which is independent and identically distributed at each pixel. We apply the same q alignment analysis to features from the last linear layer of the network before its readout layer. We found that the same general trends in alignment as described for the cortical data appear in trained, but not untrained, networks (Figure 3b). Specifically, the exemplar variance alignment with the same and other in-class variance is significantly above chance, and also larger than the alignment to the between-class axis.

**Figure 3:**
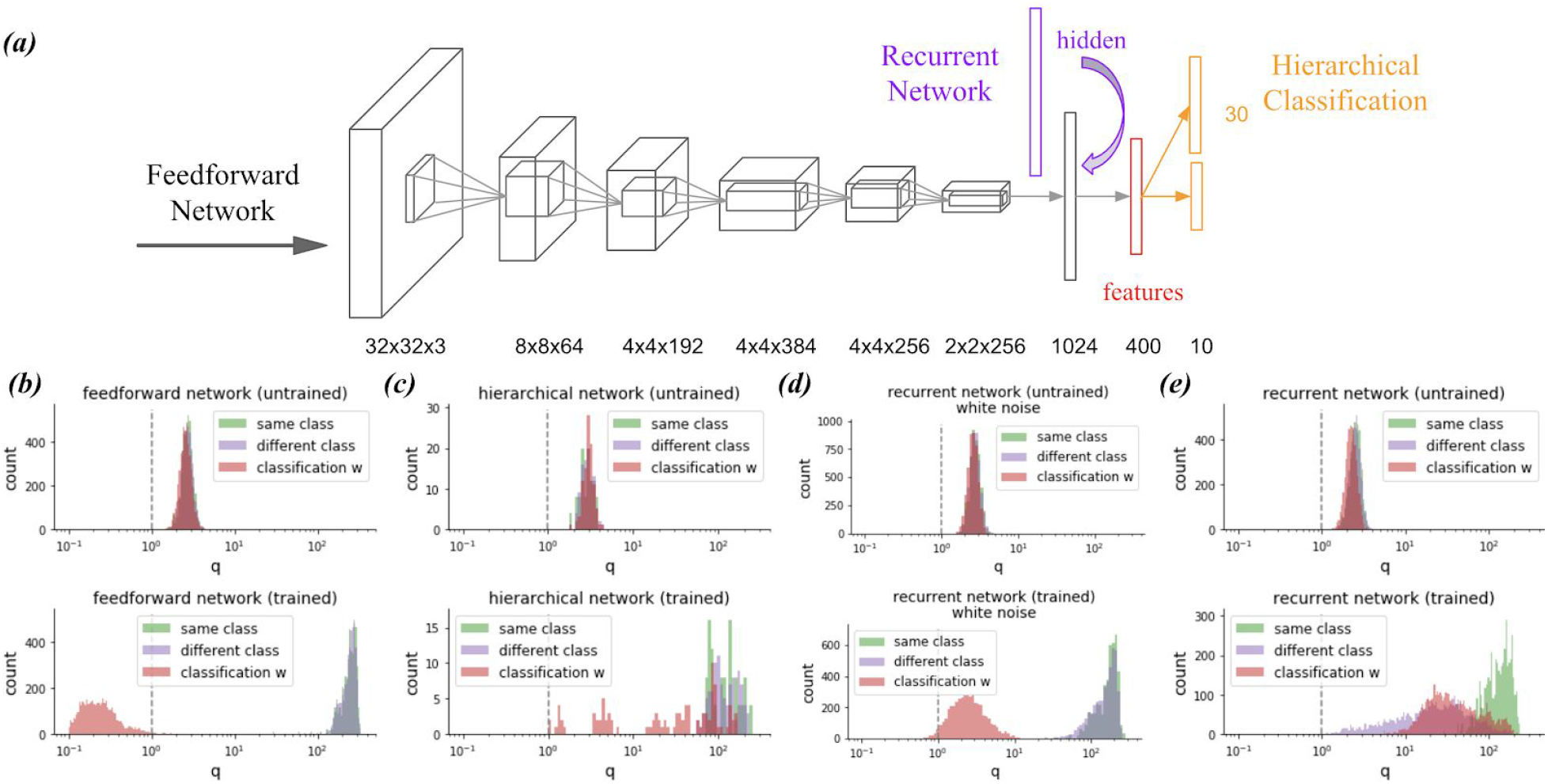
(a) Alexnet structure. (b) q for untrained and trained feedforward networks with white noise. (c) q for untrained and trained feedforward network on the hierarchical classification task. q for untrained and trained recurrent networks with white noise. (e) q for untrained and trained recurrent networks with correlated noise. For b-e, q is computed for the 1st PC.

One difference in the network statistics is that the q alignment to the classification axis w is less than 1.0. That is, the variability orthogonal to the classification axis (w) is reduced to be less than chance in the trained network (Figure 3b, bottom, red histogram). However, the q alignment to w is observed to remain larger than 1 in the neural data. We hypothesize that one of the reasons for this discrepancy is that the brain classifies inputs at multiple levels, while the trained network only needs to classify among the predefined classes.

To test this hypothesis, we train the same network on a hierarchical classification task. We subselect 5 superclasses of natural images from the CIFAR100 dataset, with 5 subclasses of images in each superclass. We then train the Alexnet network to classify inputs at both sub and super class levels (methods, III). The q alignment to the classification structure is indeed higher in the hierarchical network, supporting our hypothesis (Figure 3, c).

In addition to white noise in the external inputs, another potential origin of the aligned variability observed in the neural data is in the temporal correlation of the inputs. Unlike the artificial neural networks where the network processes discrete individual stimuli, the brain receives and processes inputs in a temporal stream. To account for this possibility, recurrent neural networks are trained to classify sequential stimuli (methods III). When variability arises from different inputs in the previous steps of the sequence, the trained, but not untrained networks exhibit the same alignment to the in-class variance as in the neural data (Figure 3d). However, the alignment to the classification boundary w is significantly smaller than observed in the hierarchical networks. On the other hand, when variability arises from white noise in individual inputs, the trained network exhibits alignment to in-class variance and classification axis w, but the alignment to the in-class variance decreases, in contrast to the large alignment similar to in-class variance observed in the neural data (Figure 3,e).

### The observed alignment is advantageous for generalization

Previous work characterizing the impact of correlated noise on stimulus coding has identified the general type of aligned noise we describe above to be information-limiting: noise spanning between 2 stimuli can reduce the accurate classification between the 2 stimuli (Averbeck et al. 2006). Here, we propose that -- in addition to this information limiting effect at the level of individual stimuli -- aligned noise can have a complementary, advantageous effect for generalization. Specifically, when only given a single stimulus from each class, aligned variability can benefit generalization by enabling classification of other novel stimuli of the same class.

To test this hypothesis, we simulate exemplar variance with different degrees of alignment to in-class variance, while keeping the overall structure of the variance unchanged by performing a rotation (methods IV). The simulated variance covers a wide range of levels of q alignment to in-class variance. Both the simulated and original data are tested for their performance on one-shot generalization, that is, the accuracy at classifying novel stimuli of two classes when only trained on a single stimulus from each class. We observe that datasets with larger q alignment show higher one-shot generalization accuracy (Figure 4a, Pearson’s correlation coefficient=0.26, p value=0.003). Additionally, consistent with the idea of information limiting correlations discussed above, the higher q alignment also leads to a decrease in classification accuracy between individual stimuli (Figure 4b, Pearson’s correlation coefficient=-0.62, p value=7e-15).

**Figure 4:**
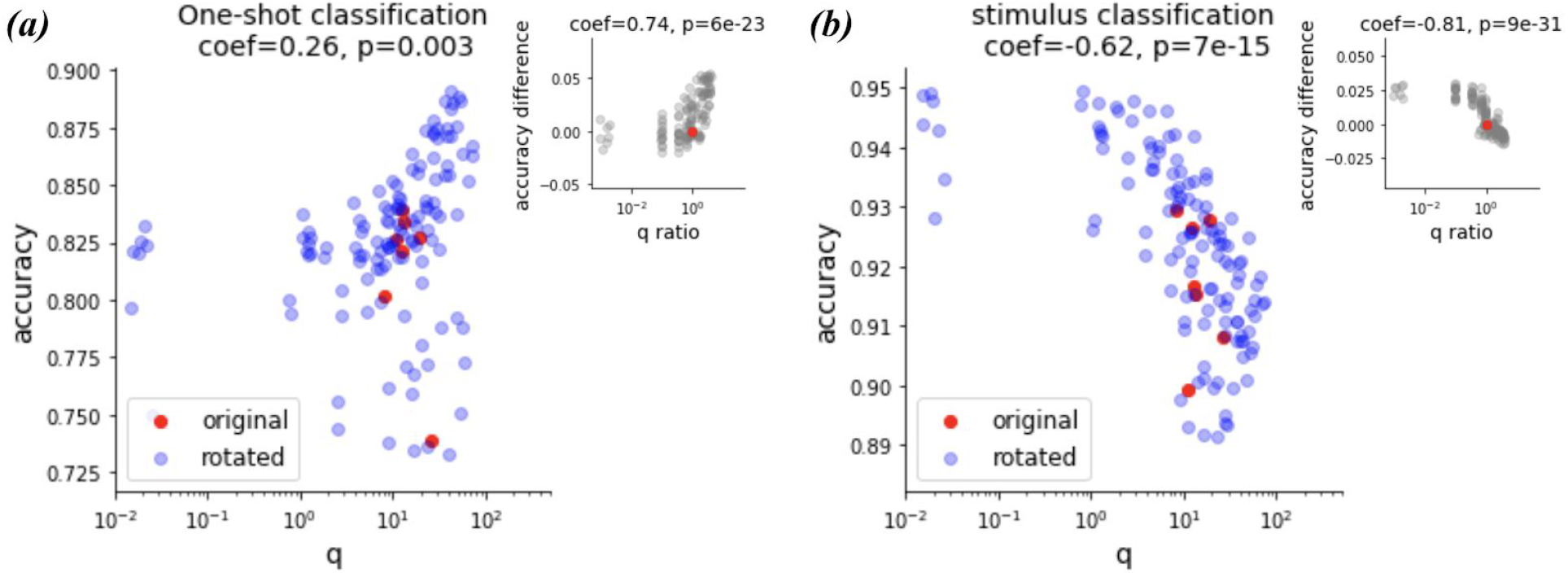
(a) one-shot generalization accuracy vs q. Insert: accuracy difference between rotated and original variability vs their corresponding q alignment to class invariant ratio. (b) accuracy at classifying individual stimulus vs q alignment to in-class variance. Insert: accuracy difference between rotated and original variability vs q ratio. In both plots, the red points represent the original neural data, the blue points represent the simulated data. The accuracy and the q alignment are averaged across all pairs of classification classes for each animal (n=7).

## Summary and Discussion

Cortical representations of sensory stimuli are very noisy. In this study we seek to characterize the geometry of variable representations of repeated stimuli in the mouse visual cortex, and to explore the potential computational roles that variability can play. We hypothesize that the role of this noise is to help generalization. To this end, we show that the variability in data from 7 mice is anisotropic, and that the variability is significantly aligned to the directions needed for generalization. We also show that such an alignment occurs in artificial neural networks trained for classification, and that it can improve one-shot learning.

Artificial neural networks trained on image classification learn to transform the representation of stimuli from pixel space -- in which a linear classifier, even with a large number of examples fails to generalize to new examples -- to a feature space, in which such a classification is possible. Functional networks to generalize in specific directions: for example, moving the image by one pixel should create the same class. Translational invariance, or asking the network to output the same class for translated images, is introduced as a prior in the architecture of the network given its convolutional structure. Other invariances, such as that within a set of features that constitute an airplane, are more difficult to express mathematically, but we rely on the learning algorithms to develop these from a set of labeled data. Using alignment measures, we observe that convolutional neural networks start with no significant differences between the alignment of exemplar and in-class variance and exemplar and between class axis. After training, these measures diverge: exemplar and in-class variance are highly aligned, while the exemplar and between class axis become orthogonal (Figure 3). Thus, injecting noise into networks gives a method for examining their structure: when one injects noise on top of an exemplar, the variation in the representation will point in the direction the network was trained to be invariant over. Even more interestingly, the exemplar variance is also aligned to in-class variance of other classes (Figure 3b), which gives the possibility for one-exemplar learning, as a learned set of in-class invariances can apply to new classes as well, enabling the new classes to be learned from one or few exemplars.

We applied the same analysis to quantify the variation in representation of scenes of natural movies in the visual cortex of mice. We use classes corresponding to continuously filmed segments in natural movies or clips; exemplars within a class are then well-separated frames within a clip. As for the case in trained neural network models, we find that: **exemplar variance is highly aligned to in-class variance, both for the same class as well as for other classes (Figure 2)**. To understand the functional consequences of the high levels of variability, we next quantified the relation between alignment and one-shot learning.

Learning from a low number of exemplars, or even a single exemplar, is a very important problem which animals often solve. For example, one can see the picture of a colleague from their website, and recognize them when they meet at a conference even if seeing them from a profile and even if the person is older than seen in the photograph. For this to work, the rules under which a new exemplar undergoing a transformation should maintain its class identity have to be generalizable from other classes (i.e., from person to person). In the case of recognizing a person from an older picture from a different angle, as humans we already understand the transformations caused by 3D rotations, as well as the features which are and are not transformed by aging. For the recordings from mouse visual cortex, we see that exemplar variance aligns with in-class variance from other classes too. The same is true for the examples in which we injected isotropic noise into the first layer of networks trained to do classification (Figure 3 b-d) but not when the injected noise was highly correlated to the previous state (Figure 3e).

To more directly test the role of alignment of exemplar variance on one-shot learning, we constructed a linear classification boundary from just two exemplars (one per class) from the mouse recordings. We find their capacity to generalize to classify examples from the same two classes to be quite good (∼80%). More importantly, when we artificially perturb this alignment, increasing the exemplar to in-class variance alignment results in an increased capacity to generalize, but at a cost of the capacity to identify individual stimuli (see Figure 4 for data averaged across mice, and Supp Fig 2 for aggregated data across mice). **From this we conclude that alignment of exemplar to in-class variance is useful for one-shot learning**.

However, there is a discrepancy between the alignment of exemplar variance to the between-class axis found in experimental data versus artificial neural network models trained on classification. In experimental data, the alignment to not only within-class variance, but also to the between-class axis, is larger than chance. We explored several possibilities for this, and found that it is reproduced in artificial neural networks trained on a hierarchical classification task. This said, we cannot exclude other computational or mechanistic explanations, which may include both recurrence and early stopping during training. We leave a detailed exploration for future work.

Finally, we note that -- while this study is focused on cortical representations found in biology -- we believe that it suggests a strategy to enhance one-shot learning with pre-learned class invariances in artificial neural networks as well: noisy versions of single exemplars can be replayed to the networks many times to obtain a representative distribution for the class. This can be viewed as a form of data augmentation (Shorten and Khoshgoftaar 2019) in which the transforms are automatically aligned to the in-class variances of the previously learned classes.

## Methods

### I. Neural Data Collection

We reused the data described in (Hu et al. 2019) for analysis. For completeness, we reproduce the text describing data collection here.

#### Electrophysiological recordings

In vivo recordings were performed in the visual cortex of awake, head-fixed mice using up to six Neuropixels probes (Jun et al. 2017). The stimulus consisted of 11 natural movie clips ranging from 1 to 6 s each for a total of 40 s were selected at random from a large database of natural movies and were repeated 200 times. All spike data were acquired with a 30-kHz sampling rate and recorded with the Open Ephys GUI (Siegle et al. 2017). A 300-Hz analog high-pass filter was present in the Neuropixels probe, and a digital 300-Hz high-pass filter (3rd-order Butterworth) was applied offline prior to spike sorting. Spike times and waveforms were automatically extracted from the raw data using Kilosort2 (Stringer et al. 2019). After filtering out units with “noise” waveforms using a random forest classifier trained on manually annotated data, all remaining units were packaged into the Neurodata Without Borders format (Teeters et al. 2015) for further analysis. This resulted in 2695 total units across seven mice.

#### Animal preparation

All experimental procedures were approved by the Allen Institute for Brain Science Institutional Animal Care and Use Committee. Five weeks prior to the experiment, mice were anesthetized with isoflurane, and a metal headframe with a 10-mm circular opening was attached to the skull with Metabond. In the same procedure, a 5-mm-diameter craniotomy and durotomy was drilled over the left visual cortex and sealed with a circular glass coverslip. Following a 2-week recovery period, a visual area map was obtained through intrinsic signal imaging (Juavinett et al. 2017). Mice with well-defined visual area maps were gradually acclimated to the experimental rig over the course of 12 habituation sessions. On the day of the experiment, the mouse was placed under light isoflurane anesthesia for 40 min to remove the glass window, which was replaced with a 0.5 mm thick plastic window with laser-cut holes (Ponoko, Inc., Oakland, CA). The space beneath the window was filled with agarose to stabilize the brain and provide a conductive path to the silver ground wire attached to the headpost. Any exposed agarose was covered with 10,000 cSt silicone oil, to prevent drying. Following a 1-2 hour recovery period, the mouse was head-fixed on the experimental rig. Up to six Neuropixels probes coated in CM-DiI were independently lowered through the holes in the plastic window and into the visual cortex at a rate of 200 µm/min using a piezo-driven microstage (New Scale Technologies, Victor, NY). When the probes reached their final depths of 2,500–3,500 µm, each probe extended through the visual cortex into the hippocampus and thalamus.

#### Data preprocessing

Three consecutive movie frames (100ms) are binned as one “analysis” frame (we will use the short term of frame for the analysis frame throughout the study) and difference frames from the same movie clips are treated as stimuli from the same class, with its corresponding neural representation the total counts of spikes during the 100ms time bin. The neural data are sub selected for time points where the animals are stationary (running speed < 5) as a control for brain states. To ensure enough stimuli per class in further analysis, only movie clips longer than 2.5 sec are included, with a total of 9 clips. The starting 3 bins of each selected movie clips are excluded from the analysis to avoid brain response not to the presented stimuli but to the movie clip transition.

### II. q alignment quantification

Given fixed stimulus i and axis w in the neural space, q alignment quantifies the degree of alignment between the exemplar variance for stimulus i to axis w:

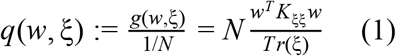

The exemplar variance is represented by ξ_*i*_∈ *R*^*T* ×*N*^, where *T* is the number of trials and *N* is the number of neurons and each row in ξ_*i*_ represents the single-trial variability (trial-average subtracted population response). 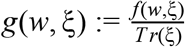 is the fraction of ξ_*i*_ variance along axis w, where *f* (*w*, δ) = *w*^*T*^ *K*_ξξ_ *w* is the variance along axis w and *Tr*(ξ), the trace of the matrix ξ, gives the total variances of ξ. Then, q is defined as the normalized fraction of variance along w, where 1/N is the expected fraction of variance along any random axis in the neural space. Therefore, a q value of 1 suggests that the variability spans in the direction *w* as much as by chance, while a larger (or less) q value suggests that the variability structure is more (or less) aligned to the chosen axis *w*.

The *q*_*multi*_ alignment quantification takes multiple axes into account and is the weighted average of the *q* alignment to different axis:

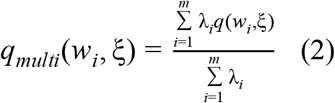

For alignment to the same/other class variance, *w*_*i*_ is the ith principal component of the trial-averaged signal for the class and λ_*i*_is its corresponding explained variance ratio. For alignment to the classification axis, *w*_*i*_ is the linear SVM classifier defined on the training data after projection to the previously defined *i* − 1 classification axes. Specifically, given the training data for the two classes *A* ∈ *R*^*k*×*N*^ (k is the number of training samples and N is the number of neurons), *w*_*i*_ is the linear SVM classifier defined on the projected data:

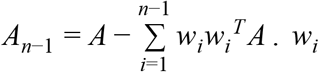 is also projected to remain orthogonal to all previously identified classification axes {*w*_1_, *w*_2_, …, *w*_*i*−1_}. The q alignment to each *w*_*i*_ is then weighted by the accuracy of *w*_*i*_ at classifying the original data *A*.

#### The Wilcoxon signed-rank test

The Wilcoxon signed-rank test is used to test whether the single-trial variability is more aligned to in-class variance of the same class, in-class variance of different classes, as well as the discrimination structures, than chance. Specifically, it tests whether the difference between the log(q) distribution and the chance level (q=1, log(q)=0) is symmetric about zero. The null hypothesis is that the log(q) distribution comes from the same distribution as q=1.

#### The Wilcoxon rank-sum test

The Wilcoxon rank-sum test is used to test that the single-trial variability is more aligned to in-class variance than to the discriminaion structure. The log(q) distribution for the two alignments are compared and the null hypothesis is that one sample is drawn from the same distribution.

### III. Artificial Neural Networks

The feedforward network adopts the Alexnet architecture and is trained on the CIFAR10 dataset to classify 10 classes of natural images. To test the variability structure for the untrained and trained networks, inputs augmented with random noise are repeatedly passed into the network. Specifically, for the feedforward network, 50 randomly selected stimuli from each of the 10 classes are repeatedly passed into the network 100 times, each time added with white noise (random noise drawn from the Gaussian distribution with zero mean and unit variance). The activity pattern at the feature layer is saved as the network activity for further alignment analysis.

The hierarchical network is trained on natural images from 25 classes, grouped into 5 superclasses (sub selected from the CIFAR100 dataset). The network adopts the Alexnet architecture, except that the final readout layer is resized to 30, with 5 units predicting labels for the 5 superclasses, and the rest 25 units predicting labels for the 25 subclasses. Therefore, the network is trained for a direct linear readout of both the class and the superclass label from the feature layer. To test the network variability structure, the averaged inputs for each subclass (averaged across stimuli in the CIFAR100 test dataset of the same subclass) is passed into the network 100 times, each time added with white noise. The activity pattern at the feature layer is saved as the network activity for further alignment analysis.

The recurrent network is trained to classify a temporal sequence of CIFAR10 inputs. Specifically, the network is trained to output the correct class label at each time point for the corresponding input. At each time point, the hidden layer activation from the previous time point is passed down and concatenated with the hidden layer of the current time point, before passed on to the feature layer for classification readout. The hidden state is initialized at zero and the training sequences are randomly selected for each training epoch. To test the network variability structure, 50 randomly selected stimuli from each of the 10 classes is passed into the network 100 times, with each stimuli preceded by 3 randomly selected stimuli from the same class. The activity pattern at the feature layer is saved as the network activity for further alignment analysis. In addition to the temporal variability, the recurrent network is also tested with white noise where the 50 stimuli are passed into the network 100 times, each time added with white noise while keeping concatenated hidden activations from the previous time point a copy of the hidden state at the current time point.

All networks are trained with the Adam optimizer (Kingma and Ba 2014). The feedforward and the recurrent networks use the CrossEntropyLoss loss function. The hierarchical network uses the BCEWithLogitsLoss loss function. All untrained network weights are initialized from a uniform distribution 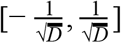, where *D* is the size of the network layer. For each of the three types of networks, 5 different networks are trained until the test accuracy plateaus.

### IV. Classification

#### Simulations of exemplar variance

To simulate exemplar variance with different q alignment to in-class variance, the variance ξ is rotated in the two-dimensional plane defined by its own variation (principal component) and the in-class variance. To rotate the exemplar variance ξ in the plane defined by two axes orthogonal *u, v* to different angles θ, ξ is multiplied by the rotation matrix *M*: δ^*rotated*^ = δ*M*.

The rotation matrix *M* is given by:

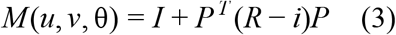

where *I* ∈ *R*^*NxN*^ and *i* ∈ *R*^2*x*2^ are the identity matrix; *R* = [[*cos*(θ),− *sin*(θ)], [*sin*(θ), *cos*(θ]], with θ the rotated angle; *P* = [*u, v*]. *u* is chosen as the same class variance (top PC), and *v* is the exemplar variance top or last PC. The exemplar variance is rotated in both planes to different angles from 0 to π, with π/10 increments to recover a wide range of q alignment.

#### One-shot generalization

For each pair of classes, one stimulus is randomly selected from each class and the single-trial representations for the two selected stimuli are used as the only training data for classification between the two classes. The trained classifier (linear SVM) is tested on the other unseen stimuli and the accuracy is averaged across all possible pairs of classes and training stimuli pairs.

#### Stimulus classification

For each pair of stimuli, we split the single-trial representations in half. The classifier is trained on half of the data and tested for accuracy on the other half. The stimulus accuracy is taken as the averaged accuracy across all stimuli pairs.

## Acknowledgments

We wish to thank the Allen Institute founder, Paul G. Allen, for his vision, encouragement and support. We wish to also thank Brian Hu, Ramakrishnan Iyer and Josh Siegle for discussions following the work presented in (Hu et al. 2019) which led to the analyses described here.

## Supplementary figures

**Supplemental Figure 1:**
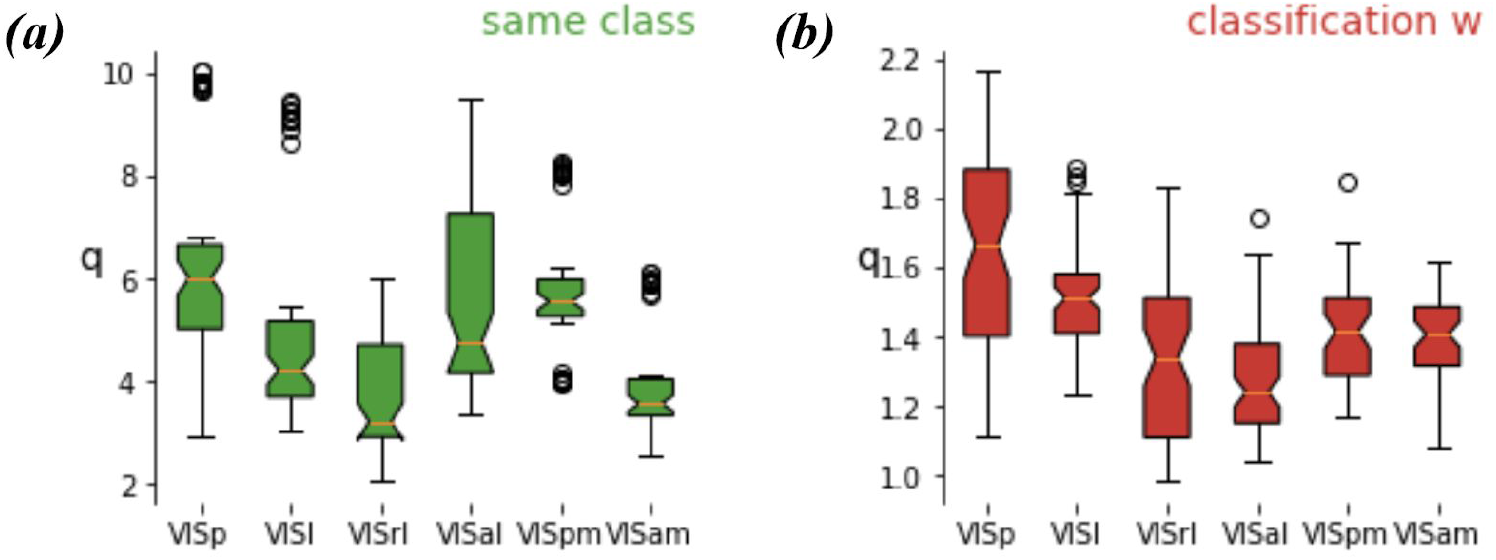
(a) q alignment to the same class variance (top 10 PCs) for different visual areas. (b) q alignment to the classification structure (top 10 axes) for different visual areas.

**Supplemental Figure 2:**
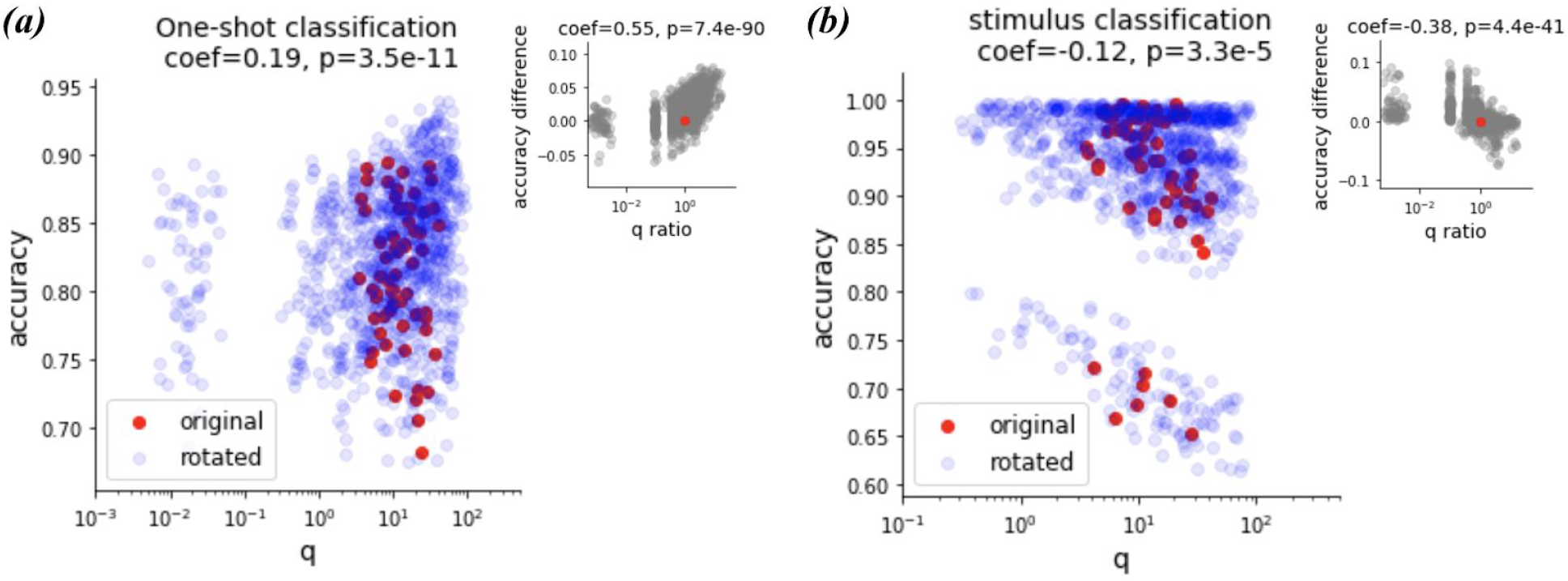
one-shot generalization accuracy vs q. Insert: accuracy difference between rotated and original variability vs their corresponding q alignment to class invariant ratio. (b) accuracy at classifying individual stimulus vs q alignment to in-class variance. Insert: accuracy difference between rotated and original variability vs q ratio. In both plots, the red points represent the original neural data, the blue points represent the simulated data. The accuracy and the q alignment are averaged across all pairs of classification classes for each class, with a total of 9 classes per animal, 7 animals (n=63).

